# Next generation whole genome sequencing of *Plasmodium falciparum* using NextSeq500 technology in India

**DOI:** 10.1101/068676

**Authors:** Alassane Mbengue, Pragya Namdev, Tarkeshwar Kumar, Kasturi Haldar, Souvik Bhattacharjee

## Abstract

*Plasmodium falciparum* is a protozoan parasite that causes the deadliest form of human malaria. Although, malaria burdens worldwide have decreased substantially over the last decade (WHO, 2014), genetic variation and adaptation by parasite strains against drugs and vaccines present significant challenges for the elimination of malaria (Ariey et al., 2014; Neafsey et al., 2015). India has formally launched a malaria elimination campaign (NVBDCP, 2016). Therefore, early in-country detection of drug resistance and/or immune evasion will be important for the program. Presently, the majority of surveillance methods in India detect a limited number of known polymorphisms (Campino et al., 2011; Chatterjee et al., 2016; Daniels et al., 2008; Mishra et al., 2015; Neafsey et al., 2012; Neafsey et al., 2008). A recently reported amplicon sequencing method enables targeted re-sequencing of a panel of genes (Rao et al., 2016). However, the capacity to identify new genes of resistance/immune evasion by whole genome sequencing (WGS) through next generation sequencing (NGS) in India, has remained elusive. Here we report the first WGS of *P. falciparum* strain performed by Eurofins Genomics India Pvt. Ltd at its Bengaluru division within 40 days of sample submission. Our data establish that timely, commercial WGS through NGS in India can be applied to *P. falciparum* to greatly empower the malaria elimination agenda in India.

---

The malaria burden has reduced significantly in India in last 15 years, giving impetus to the development of the *National Framework for Malaria Elimination in India* 2016–2030 (NFME, 2016). However, it is recognized that many of the achievements are fragile and need vigilant support. In the face of parasite resistance to all known antimalarials at a global level (WHO, 2015), emergence of drug resistant parasites needs to be closely monitored to systematically evaluate efficacy of therapies in India. Further, as vaccination strategies mature, their administration may trigger different adaptive responses in areas with vastly different levels of host immunity associated with a wide range of parasite transmission densities prevalent in India (see Figure. 1).

**Figure 1.**
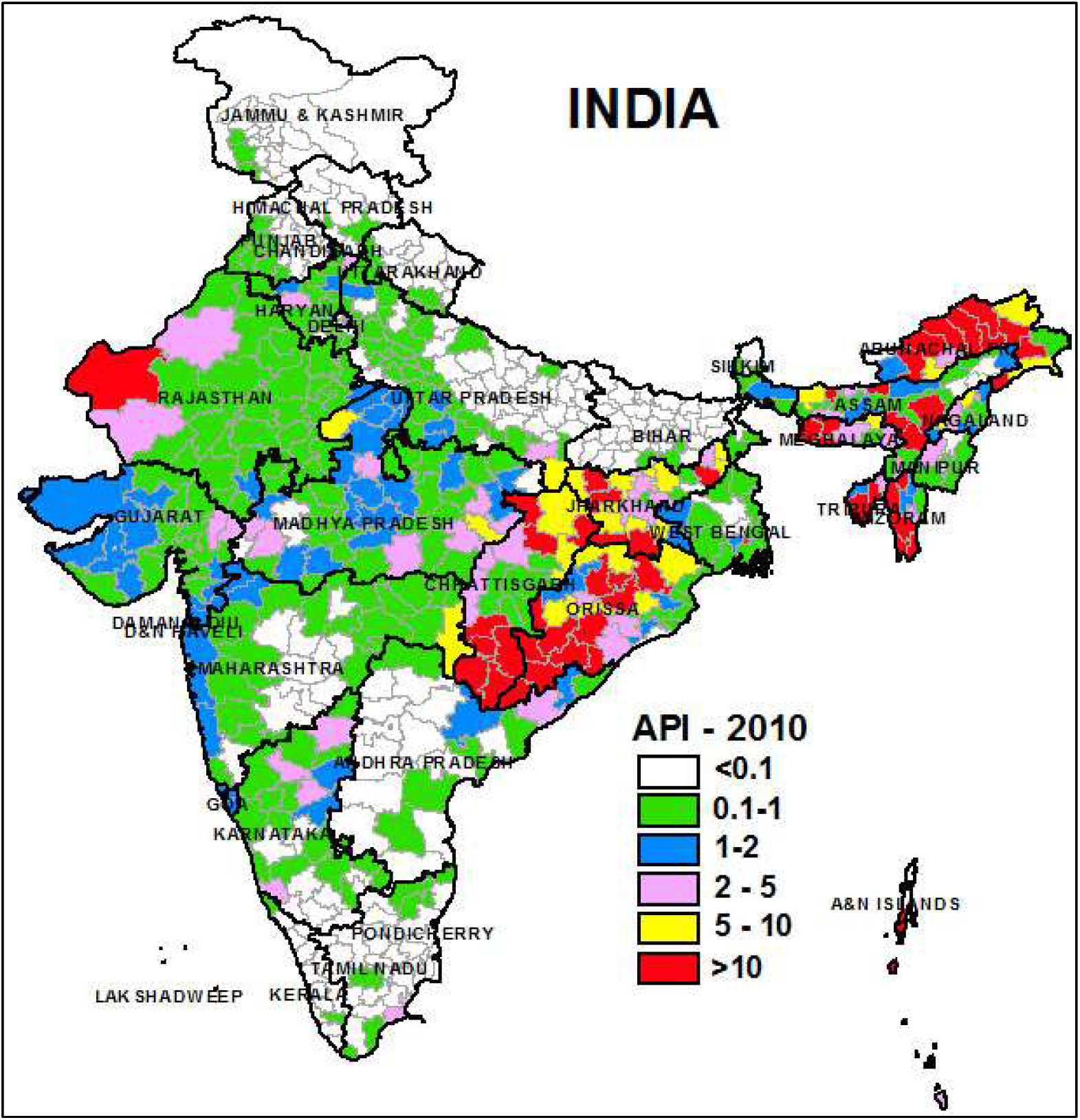
Distribution of malaria in India. Schematic representation showing malaria distribution in India during 2015, based on annual parasite index (API). Adapted from http://www.nvbdcp.gov.in/maps.htm.

Therefore, understanding parasite genetic backgrounds and genetic diversity is key to the elimination agenda. It is critical to monitor parasite strain dynamics and virulence, residual pools of parasites, transmission profiles emergence of resistance to therapeutic strategies employed in continued reduction and elimination. Genomic technologies are powerful in elucidating genetic backgrounds. Methodologies adapted for malaria-endemic countries increasingly provide sequencing support for identification of known single nucleotide polymorphisms (SNPs) or new polymorphisms in one or more genes to monitor drug or immune selection (Bailey et al., 2012; Chang et al., 2012; Galinsky et al., 2015; Gandhi et al., 2012; Manske et al., 2012). Recently a *P. falciparum* mitochondrial genome was sequenced through Sanger sequencing (Tyagi et al., 2014). However, WGS needed to completely map the genetic template is not yet widely accessible in India.

WGS is particularly important in monitoring hitherto unknown resistance markers and immune adaptation that cannot be comprehensively deciphered from analyses of a defined set of parasite genes. In context of drugs, resistance to artemisinins, frontline antimalarials to which we still have no replacements, is thought to likely involve multiple genetic loci, in addition to the major determinant *pfkelch13* (Ariey et al., 2014; Miotto et al., 2015). Further *pfkelch13* mutations confer different levels of resistance in different strain backgrounds (Straimer et al., 2015) emphasizing the need for whole genome sequencing to rapidly understand and distinguish strains in efforts to track and contain resistance before it proceeds to relapsed, difficult-to-treat infections.

WGS of Indian isolates has lagged behind in analyses of parasite strains from other endemic regions of the globe including Africa, China, SE Asia, South and-Central America (Bailey et al., 2012; Borrmann et al., 2013; Chang et al., 2012; Cheeseman et al., 2012; Daniels et al., 2015; Obaldia et al., 2015; Tan et al., 2012). This could be greatly overcome by in-country capacity for WGS in India. To overcome this bottle-neck we undertook WGS and associated bioinformatics analyses of *P. falciparum* strain 3D7 with a commercial vendor Eurofins India Pvt. Ltd in Bengaluru. We present this as a first report of WGS of *P. falciparum* in India.

## Materials and Methods

### Maintenance of P. falciparum culture

*P. falciparum* 3D7 parasites were propagated in A+ human erythrocytes in RPMI supplemented with 0.5% Albumax II, 0.2 mM Hypoxanthine, 11 mM Glucose, 0.17% NaHCO_3_, 10 µg/ml gentamycin and maintained at 37°C in 5% CO_2_ in a humidified incubator, as described earlier (Bhattacharjee et al., 2012). Cultures were monitored daily by Giemsa staining of methanol-fixed smears and fed as necessary. Parasitemia were usually maintained below 10% for healthy culture growth.

### Preparation of P. falciparum 3D7 genomic DNA (gDNA)

Total genomic DNA was isolated from P. falciparum 3D7 infected red blood cells using Quick-gDNA miniprep kit from Zymo Research (Cat # D3024) following manufacturer instructions. The gDNA was eluted from column using DNase, RNase-free water and stored at -20°C until shipment to Eurofins Genomics India Pvt. Ltd.

### Shipment of gDNA from the laboratory to the WGS sequencing facility at Whitefield, Bengaluru, India

The *P. falciparum* 3D7 genomic DNA sample was shipped to Eurofins Genomics India Pvt. Ltd. in dry ice by courier.

### Qualitative and quantitative analysis of gDNA

The quality of the gDNA was assessed by resolving in 0.8% (w/v) agarose gel at 120 V for 30 min. For quantitative estimation, 1 µl of gDNA sample was used to determine the A260/280 ratio using Nano-drop One (Thermo Fisher Scientific) and concentration was measured using Qubit^®^ 3.0 Fluorometer (Thermo Fisher Scientific).

### Preparation of 2 x 150NextSeq 500 library

The paired-end sequencing library was prepared using NEBNext Ultra DNA Library Prep Kit (NEB). Briefly, approximately 1µg of gDNA was fragmented by Covaris M220 ultrasonicator to generate a mean fragment distribution of 400 bp. Covaris shearing generates dsDNA fragments with 3’ or 5’ overhangs. The fragments were then converted to blunt ends using End Repair Mix. The 3’→5’ exonuclease activity of this mix removes the 3’ overhangs and the 5’→3’ polymerase activity fills in the 5’ overhangs. This was followed by adapter ligation to the fragments. This strategy ensures a low rate of chimera (concatenated template) formation. The ligated products were size selected using AMPure XP beads (Beckman Coulter). The size-selected products were PCR amplified with the index primers according to manufacturer instructions.

### Quantity and quality check (QC) of library on Tape Station

The PCR amplified libraries were analyzed in TapeStation 4200 (Agilent Technologies) using High Sensitivity (HS) D5000 Screen Tape assay kit as per manufacturer instructions.

### Cluster Generation and Sequencing

After obtaining the Qubit concentration for the libraries and the mean peak size from Agilent TapeStation profile, the PE Illumina library was loaded onto NextSeq 500 for cluster generation and sequencing. Paired-End sequencing allowed the template fragments to be sequenced in both the forward and reverse directions on NextSeq 500. The reagents provided in the kit were used in binding of samples to complementary adapter oligos on paired-end flow cell. The adapters were designed to allow selective cleavage of the forward strands after re-synthesis of the reverse strand during sequencing. The copied reverse strand was then used to sequence from the opposite end of the fragment.

### Sequencing data analysis

NGS of P. falciparum 3D7 PE library was performed by Eurofins Genomics India Pvt. Ltd. on an on NextSeq 500 using 2 × 150 bp chemistry. This generated a total number of 34,399,710 raw reads with 10,331,446,863 bp and 10.3 GB of raw data. These raw reads were filtered using Trimmomatic v0.35 with the following stringent parameters: (i) adapter trimming; (ii) SLIDING WINDOW- trimming of 20 bp, cutting once if the average quality fell below a threshold of 20; (iii) LEADING- cutting bases at the start of a read if below a threshold quality of 30; (iv) TRAILING- cutting bases at the end of a read if below a threshold quality of 25; and (v) MINLENGTH-dropping a read if >100 bp. After filtration, the total number of reads and bases were 23,951,749 and 7,050,844,590, respectively. The final data size at the end of this step was 7 GB. The cumulative data were aligned against reference *P. falciparum* 3D7 genome sequence, downloaded from NCBI (ftp://ftp.ncbi.nlm.nih.gov/genomes/all/GCF_000002765.3_ASM276v1/GCF_000002765.3_ASM276v1_genomic.fna.gz). For confirmation, we assessed the sequencing coverage of two drug-resistance genes: *pfkelch13* (PlasmoDB id PF3D7_1343700), pfmdr1 (PlasmoDB id PF3D7_0523000) associated with artemisinin resistance.

## Results and Discussion

### Quality control on agarose gel

The *P. falciparum* 3D7 genomic DNA was quantitated using Qubit 3.0 Fluorometer (Thermo Fisher Scientific). The estimated concentration and amount was 93.5 ng/µl and 1.87 µg, respectively. The quality check of the genomic DNA in 0.8% agarose gel revealed high molecular weight band characteristic of intact genomic DNA. DNA smearing, representing genomic DNA fragmentation during purification was minimally seen (Figure 2).

**Figure 2.**
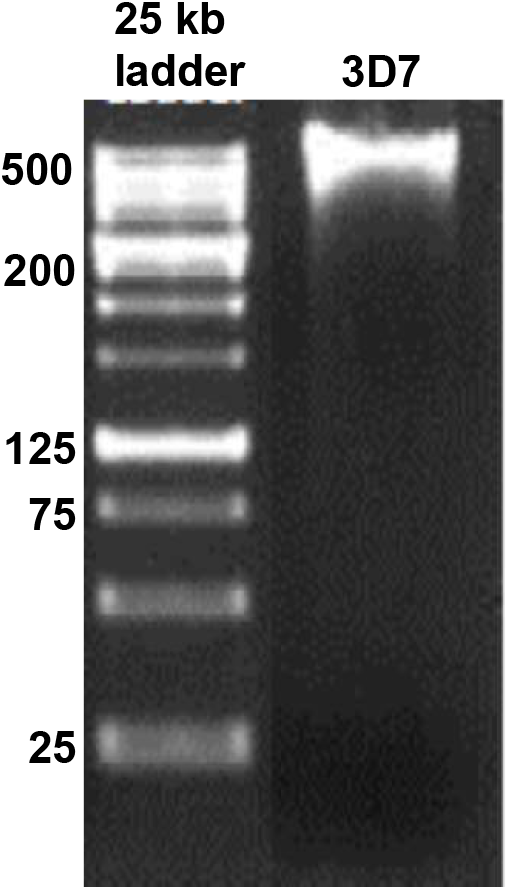
Quality check of P. falciparum 3D7 genomic DNA. Total genomic DNA from the laboratory-adapted *P. falciparum* 3D7 culture was resolved using 0.8% agarose gel. Standard DNA 25 kb ladder is shown at the left.

### Library profile in Agilent Tape station

Following quality check, the paired-end (PE) libraries were prepared using NEB Next Ultra DNA Library Prep Kit for Illumina using manufacturer instructions. The libraries were sequenced on NextSeq 500 using 2 × 150bp chemistry. The library profile is shown in Figure 3.

**Figure 3.**
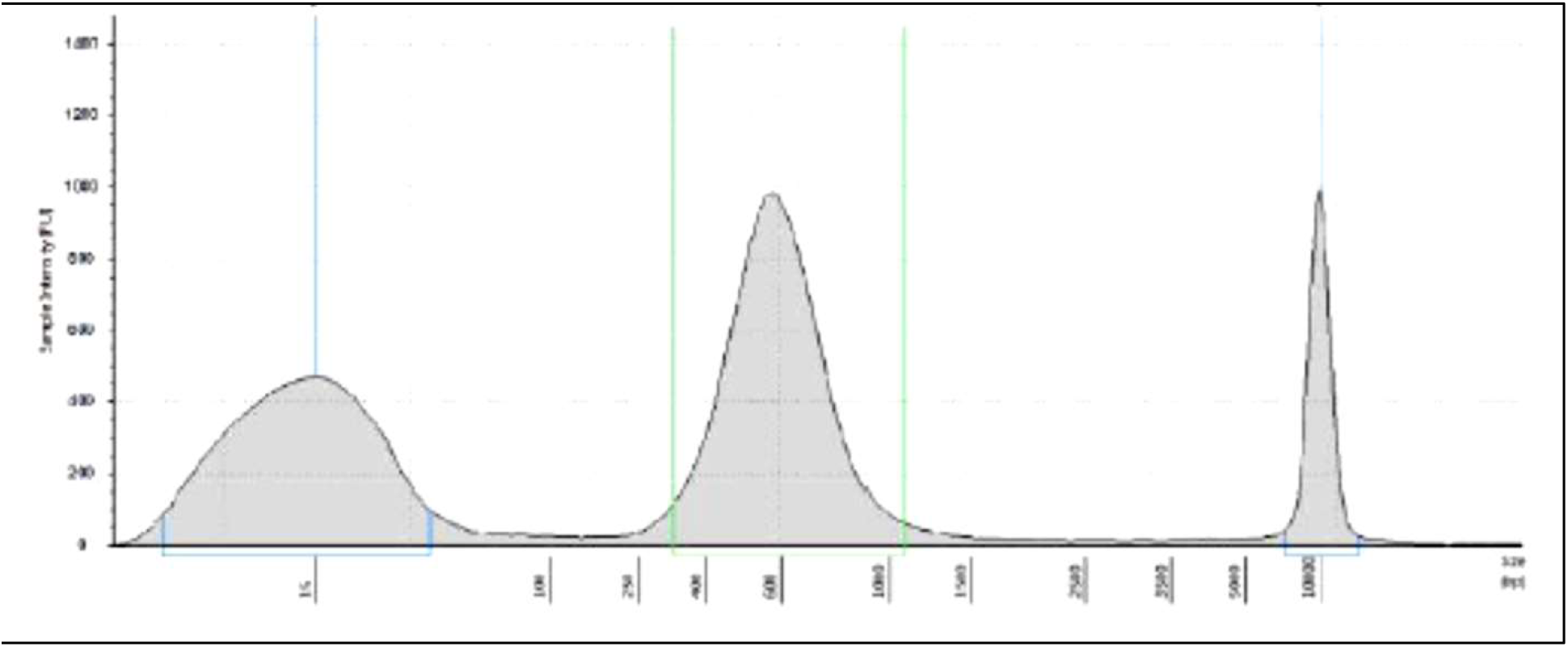
Library profile of sample 3D7 on Agilent Tape Station HS D5000 screen tape. Highlighted between green perpendicular line shows the profile from 319-1080 bp with an average size of 598 bp. This represents 89.9% of the total library at a concentration of 780 pg/µl (2170 pmol/l).

### Raw and filtered data statistics

The total number of reads and bases from the 3D7 sample was 34,399,710 and 10,331,446,863 bp, respectively. This generated 10.3 GB of raw data. After filtration, the total number of reads and bases were 23,951,749 and 7,050,844,590 bp. The final data size was 7 GB.

### Mapping reads to the reference genome

The *P. falciparum* 3D7 genome, downloaded from ftp://ftp.ncbi.nlm.nih.gov/genomes/all/GCF_000002765.3_ASM276v1/GCF_000002765.3_ASM276v1_genomic.fna.gz, was used as a reference genome. High quality reads were mapped against this reference genome with size of 23.27 Mb using BWA mem (*version 0.7.12-r1039*) with default parameters including minimum seed length of 19 and penalty of mismatch at 4.

Mapping was performed by indexing of the reference genome and aligning filtered reads to the reference index. The alignment was obtained in BAM file format, which was further used to generate consensus genome for 3D7 sample. Overall, the mapping revealed 22,255,033 reads with a mapping percentage of 93%. The total number of unmapped reads was 1,696,716 and the genome coverage was 99.99%. The bioinformatics analysis workflow is shown in Figure 4.

**Figure 4.**
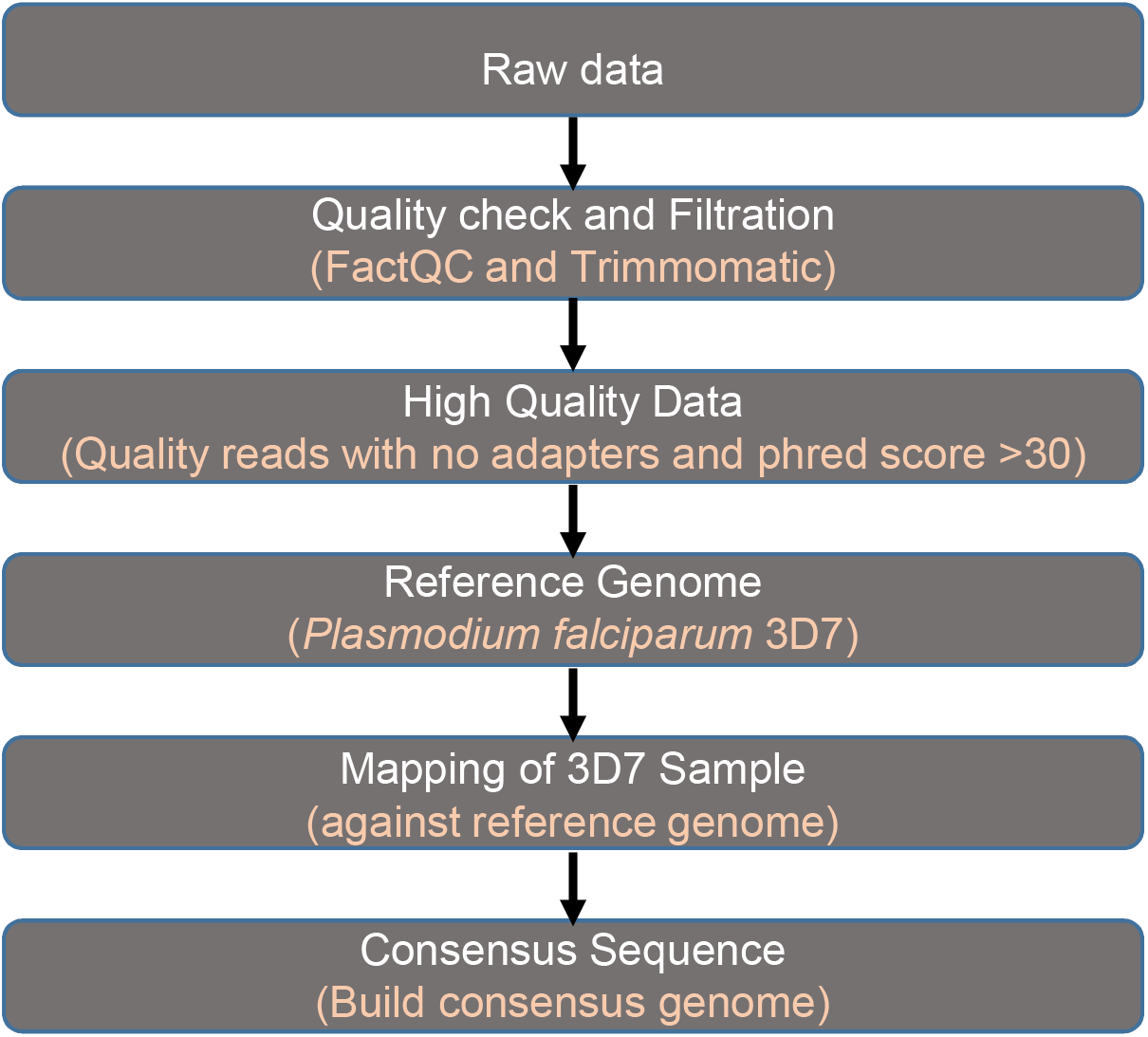
Bioinformatics analysis workflow. Raw data reads from the sequenced library were filtered and mapped against the reference *P. falciparum* 3D7 genome from NCBI (ftp://ftp.ncbi.nlm.nih.gov/genomes/all/GCF_000002765.3_ASM276v1/GCF_000002765.3_ASM276v1_genomic.fna.gz) with set default parameters, as described in Materials and Methods.

## Concluding Comments

Our work adds WGS to the genomic tool kit for studies of *P. falciparum* strains in India. Its immediate application to recently isolated Indian strains that have been cultured in the laboratory will yield unbiased genetic information of *P. falciparum* strains in India (John White III, 2016). Since these represent ∼ 50% of collected isolates, they are expected to yield comprehensive regional strain analyses which are currently missing to fully understand emergence of artemisinin resistance and potentially future vaccine trials. As other aspects of laboratory and field capacities are further strengthened in-country, WGS will allow definitive strain analysis at the national level needed for malaria elimination in India.

